# Cervical collagen network porosity assessed by SHG endomicroscopy distinguishes preterm and normal pregnancy – a pilot study

**DOI:** 10.1101/2024.03.12.584664

**Authors:** Wenxuan Liang, Yuehan Liu, Honghua Guan, Vorada Sakulsaengprapha, Katherine Luby-Phelps, Mala Mahendroo, Xingde Li

## Abstract

**Background:** Preterm birth (PTB) is a global public health issue affecting millions of newborns every year. Orchestrated remodeling of the cervix is essential for normal pregnancy and birth, while PTB is closely related with premature cervical ripening and loss of cervical mechanical strength. The structure and organization of fibrillar collagen in the extracellular matrix are of vital importance to the biomechanical properties of the cervix. Second harmonic generation (SHG) microscopy has proved capable of revealing the progressive changes in cervical collagen morphology over the course of pregnancy. To translate this promising imaging technology to clinical practice, a flexible SHG endomicroscope has long been envisaged for label-free, non-invasive visualization of cervical collagen architecture and for assessment of PTB risk.

**Objective:** To evaluate the potential of our newly-developed SHG endomicroscope for imaging-based differentiation of cervical collagen architecture between normal pregnant mice and RU486/mifepristone-induced PTB mouse models.

**Study Design:** We undertook endomicroscopy SHG imaging of cervical collagen on two types of *ex vivo* samples: 1) frozen cervical tissue sections (∼50 µm thick) and 2) resected intact cervices, and performed SHG-image-based quantitative collagen morphology analysis to distinguish RU486 mouse models from normal pregnant mice.

**Results:** Endomicroscopic SHG images of cervical tissue sections from mifepristone-treated mouse models exhibit statistically larger collagen fiber diameter, increased pore size, and reduced pore numbers than those of normal pregnant mice. Similar changes are also observed on SHG images of subepithelial collagen fibers acquired from intact cervices by the endomicroscope.

**Conclusion:** The experiment results demonstrated that SHG endomicroscopy along with quantitative image analysis holds promising potential for clinical assessment of cervical collagen remodeling and preterm birth risk.

## 1 Introduction

Pre-term birth (PTB) is a global public health problem, affecting 12.7 million new-born babies worldwide every year, and one of every eight births in United States^1,2^. Surviving premature infants still suffer from higher rates of learning disabilities and various chronic diseases that can last a lifetime, which further present tremendous psychological and economic burden to the family and the entire society^3^. To reduce the incidence of PTB, deeper insight into the underlying mechanisms of both normal and premature birth, and reliable diagnosis technology for PTB risk assessment are of paramount importance.

It is well-known that throughout gestation, the cervix goes through dramatic remodeling, evolving from being stiff and closed to pliable enough and open sufficiently to allow the passage of a mature fetus^4^. The cervix stroma contains a rich extracellular matrix (ECM) that include type I and III collagen fibers, elastic fibers proteoglycans, and hyaluronan. Current evidence supports the programmed change in cervical compliance as regulated by a continuous turnover of collagen which ensures in pregnancy the replacement of strong, crosslinked collagen with poorly crosslinked collagen^5-7^ and increase in collagen solubility^8^, although the total collagen content stays constant^8,9^. This emphasizes that it is the morphology, architecture and other properties of cervical collagen, rather than abundance of collagen, that determine the biomechanical properties of the cervix^10,11^. Moreover, the progressive cervical remodeling process is well-conserved between human and mouse^6,12,13^.

Second harmonic generation (SHG) microscopy, with inherent depth-resolving capability and deep tissue penetration, is an appealing imaging technology for interrogating micro-structure of type I collagen in a noninvasive and label-free fashion^14-17^. Previous SHG microscopy studies have revealed the progressive changes in morphology of cervical collagen over the course of normal pregnancy, essentially a transition from straight thin fibers to kinked thickened fibers with increased inter-fiber spacing (porosity)^18^. Although the molecular origin of such morphological changes remains to be determined, the information obtained by SHG imaging still promises a valuable diagnostic tool for detecting premature or abnormal cervical ripening. To translate the power of SHG microscopy for clinical benefits and to further the investigation into cervical collagen remodeling directly on intact cervical stroma *in vivo* and *in situ*, an SHG endomicroscope is highly desired. Our group has pioneered the development of high-resolution high-performance fiber-optic flexible nonlinear endomicroscopes^19-21^. Through a series of technological innovations on background suppression and collection efficiency enhancement, our newly-developed nonlinear endomicroscope has achieved unprecedented imaging signal-to-noise ratio (SNR), enabling two-photon fluorescence and SHG endomicroscopic imaging of various label-free biological tissues with image quality rivaling a bench-top microscope^22^. Preliminary imaging study of mouse cervical tissue sections has confirmed the endomicroscope is capable of visualizing thickening of cervical collagen fibers over the course of normal pregnancy^21^; however, how well the SHG endomicroscope lends itself to capturing abnormal morphology and organization of cervical collagen remains to be investigated.

In this study, we utilize a mouse model of preterm birth to advance the applicability of SHG endomicroscopy for PTB detection. Mice treated with the progesterone receptor antagonist, mifepristone/RU486 undergo preterm birth due to the premature loss of progesterone function^23^. Similar to mice, administration of mifepristone to pregnant women accelerates cervical remodeling^24^. Images were obtained from frozen cervical tissue sections harvested from gestation day 15 pregnant mice +/-RU486 exposure for 12 hours^25^. Image analysis is undertaken, with both standard Fourier-domain methods and a novel porosity analysis algorithm specially tailored for our endomicroscopy images, to characterize altered morphology and organization of cervical collagen in the RU486 PTB model, and to distinguish them from normal pregnant mice at the same gestation time point. We then further demonstrate that our SHG endomicroscope can image subepithelial collagen directly through epithelia of intact mouse cervices and reveal similar collagen architecture features as seen in sliced tissue sections, opening up new possibilities towards *in vivo* investigation of cervical collagen morphology without the need of tissue resection.

## 2 Materials and Methods

### 2.1 SHG endomicroscopy system

Illustrated in Fig. 1A is the schematic of the SHG endomicroscopy system, whose design and working principles have been detailed in previous reports^19,22,26^. Briefly, the scanning head of the endomicroscope (purple dashed box in Fig. 1A) is constructed around a single piece of customized double-clad fiber (DCF). The excitation laser pulses of 890 nm central wavelength are delivered through the single-mode core of the DCF, and focused onto the cervical tissue by a customized achromatic miniature objective with up to 0.8 NA and 200 µm working distance (WD) in water. SHG signal photons, originating predominantly from type I collagen fibers and backscattered by the tissue, are epi-collected effectively by the same objective into the DCF, mainly captured by the large inner clad (185 µm in diameter; acceptance NA ∼0.35), and then guided back to the proximal end for detection.

**Figure 1.**
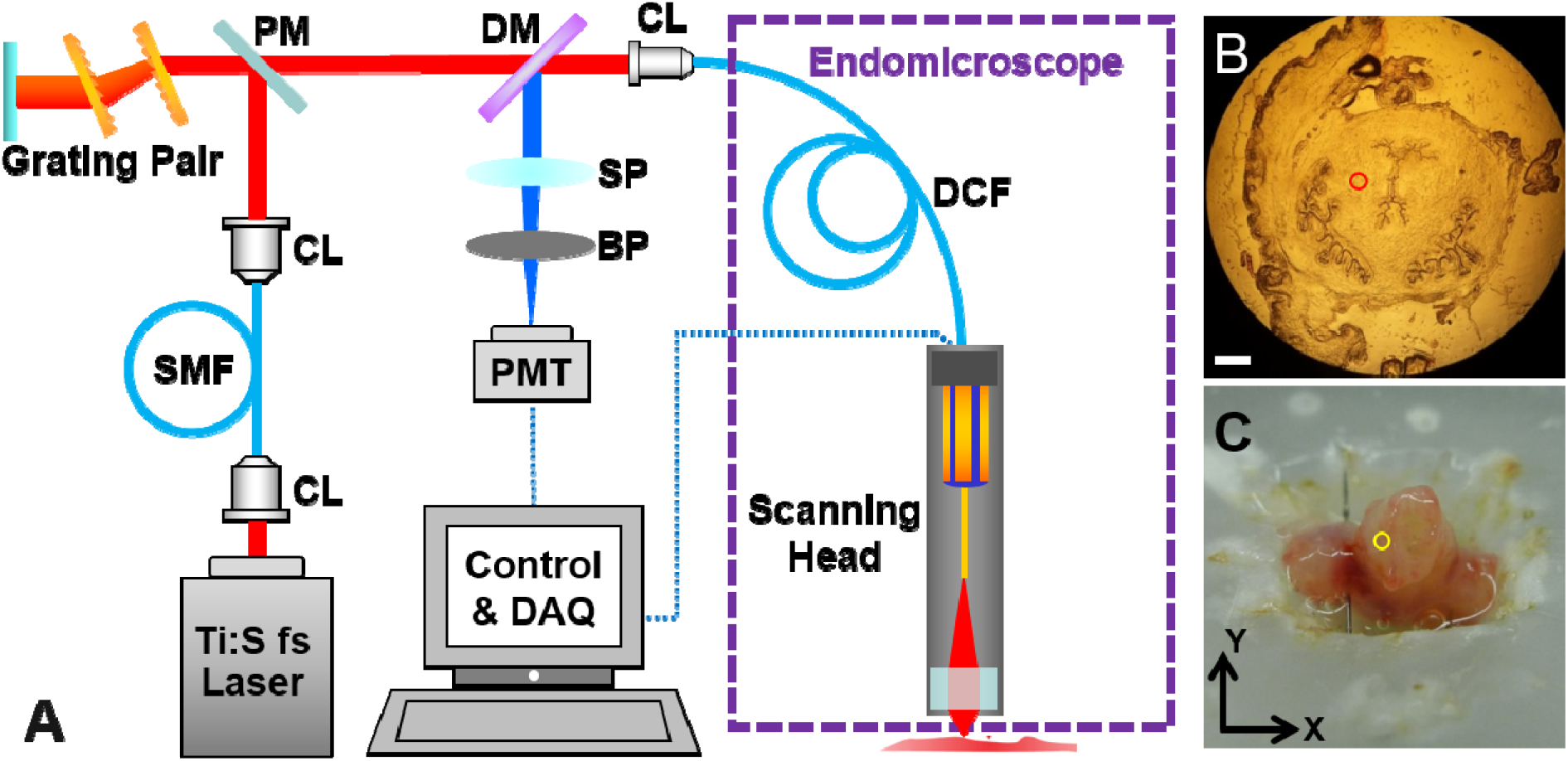
Endomicroscopy system schematic and experimental setup. (A) Optical setup of the SHG endomicroscopy system. BP: bandpass filter; CL: coupling lens or collimating lens; DCF: double clad fiber; DM: dichroic mirror; PM: pickup mirror; PMT: photomultiplier tube; SMF: single-mode fiber. (B) Wide-field microscopy photo of an example frozen cervical tissue section. Scale bar: 400 μm. (C) Photograph of an intact mouse cervix to demonstrate the imaging setup for intact mouse cervices. An example imaging field of view (or imaging spot) is illustrated by a red and yellow circle in subfigure (B) and (C), respectively.

Femtosecond laser pulses propagating through the DCF core suffer from pulsewidth broadening caused by both linear temporal dispersion, i.e. group velocity dispersion (GDD), and nonlinear spectral narrowing effect due to self-phase modulation (SPM)^27-29^. To counteract both adverse effects, we adopted a dual-fiber strategy, as shown in Fig. 1A. Near-infrared laser pulses from the Ti:sapphire laser (Chameleon Vision II, Coherent), with temporal pulsewidth of ∼150 fs, are first coupled into a piece of single-mode fiber (SMF, PM780-HP, Thorlabs); then the pulses propagate through a grating pair (600 lines/mm, Wasatch Photonics) before getting coupled into the DCF core of the endomicroscope. The separation of the grating pair is tuned meticulously to just compensate the total GDD induced by the SMF (∼25 cm long) and the DCF (∼75 cm long) together. In this way, the stretched pulse out of SMF is essentially flipped in space, and the following propagation in DCF acts essentially as a time-reversal process, canceling both linear and nonlinear effects occurred in SMF and effectively mitigating the total pulsewidth broadening^30,31^. Compared with the common dispersion compensation scheme using only the grating pair, which led to final full-width-at-half-maximum (FWHM) pulsewidth on the order of ∼400-500 fs^21,32^, such dual-fiber strategy restores the final out-of-DCF pulsewidth back to ∼150 fs, thus promoting the two-photon excitation efficiency by ∼2-3 times.

To perform lateral beam scanning, the DCF is threaded through and anchored concentrically to a small piezoelectric (PZT) tube, with an extra piece (∼11 mm long) projecting out as a freely-standing end cantilever. The surface of the PZT tube is divided into four quadrants, forming two orthogonal pairs of electrodes. By driving the two pairs of quadrant electrodes with amplitude-modulated sine and cosine waveforms (i.e. 90 degrees out of phase) respectively, and setting the frequency close to mechanical resonant frequency (typically ∼1.3 kHz) decided by the fiber cantilever, the fiber tip can be scanned into a spiral trace that covers a circular field of view (FOV)^19,26^. In this study, each raw frame consisted of 512 spirals, corresponding to a frame acquisition time of ∼0.39 s. The spatial resolution of the endomicroscope was measured to be ∼0.7 µm laterally and ∼6.5 µm axially.

The SHG emission photons guided back to the proximal end of the endomicroscope are first separated from the excitation light by a long-pass dichroic beamsplitter (FF665-Di02-25x36, Semrock), and then pass a short-pass optical filter (FF01-680/SP-25, Semrock) to further eliminate residual excitation photons. The SHG signal photons are purified by a band-pass optical filter (FF01-445/20-25, Semrock) placed in front of the photomultiplier (H10771P-40, Hamamatsu), defining a final detection band of 435-455 nm.

### 2.2 Tissue samples preparation

Normal pregnant mice (C57B16/129S) were given 0.5 mg of mifepristone (RU486) in Triolene oil (Sigma), or vehicle control (Triolene oil only) via subcutaneous injection on the morning of gestation day 15, and then sacrificed 12 hours later. Intact cervices were harvested, embedded in the optimal cutting temperature (OCT) compound (Tissue Tek, Elkhart, Indiana) and snap frozen in liquid nitrogen. For tissue section imaging, the entire cervix was further sliced transversely into 50 µm-thick sections, and mounted onto microscope slides. A microscopy photograph of an example frozen tissue section is shown in Fig. 1B.

Before imaging, samples were immersed in an appropriate amount of 0.1 M phosphate buffered saline (PBS), typically one drop for each tissue section and a petri dish full of PBS for intact cervices, to thaw in room temperature for ∼15 minutes. Then the tissue section was directly moved to the SHG endomicroscopy system and imaged without coverslip, while PBS immersion was maintained throughout imaging process. An intact mouse cervix, once thawed, was first pinned onto a home-made wax block and oriented with the ectocervix facing upwards, as shown in Fig. 1C. Then the endomicroscope was positioned gently against the ectocervical surface (without breaking the epithelial), and PBS was dripped into the space between the endomicroscope and tissue surface to maintain water immersion for the endomicroscope during imaging. Such setup, although *ex vivo*, mimics closely the *in vivo* scenario, where an endomicroscope can be maneuvered to press gently against the ectocervix for image acquisition without disturbing the endocervix or uterus. All imaging studies reported in this manuscript were conducted in compliance with the standards of humane animal care described in the National Institutes of Health Guide for the Care and Use of Laboratory Animals, using protocols approved by institutional animal care and research advisory committees at UT Southwestern Medical Center, Dallas, TX, and at Johns Hopkins University, Baltimore, MD.

### 2.3 Quantification of collagen fiber diameter

To quantify the average fiber diameter in a given SHG image, we adopted the Fourier domain method based on the two-dimensional (2D) autocorrelation function (ACF)^18,33^. The 2D ACF of each SHG image (734 × 734 pixels), denoted as ACF(*ξ, η*), is computed via the inverse 2D Fourier transform of its power spectrum (with the power spectrum defined as the modulus squared of each image’s 2D Fourier transform); here all (inverse) Fourier transform are performed via fast Fourier transform (FFT) in MATLAB (The MathWorks, Inc., Natick, Massachusetts). The central circular region 64 pixel (corresponding to ∼8.7 μm) in radius of the 2D ACF is extracted (with the origin ACF(0,0) excluded since it contains spurious peak contributed by shot noise), and then fitted to a 2D Gaussian function 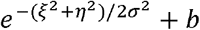 where *a* is the extrapolated amplitude of the ACF, *b* the direct current (DC) offset, and *σ* the common standard deviation for both *ξ* and *η* axes. Two times the standard deviation, i.e.2*σ*, is used as a scale to characterize fiber diameter^18^.

### 2.4 Superpixel-based image segmentation and porosity analysis

To better appreciate the connectivity of collagen fibers in the SHG images acquired by our endomicroscope, we specially designed a novel porosity analysis algorithm, which involves 1) superpixel aggregation; 2) graph-cuts pore segmentation; and 3) porosity analysis. Our algorithm starts with k-means clustering raw pixels in the SHG image, based on their grey value similarity and spatial proximity, into superpixels that are uniform in size, using the *simple linear iterative clustering* (SLIC) algorithm34,35. This clustering procedure aggregates nearby similar raw pixels into one superpixel, so as to: 1) minimize the influence of local pixel-to-pixel brightness fluctuations on segmentation, 2) capture the architecture information of collagen fibers on a more macroscopic level, and 3) reduce the computational cost as the number of pixels to handle decreases from ∼0.54 million (734×734 reconstructed from 512 spirals) to average grey value of all constituent raw pixels. Throughout this manuscript, we will denote the i-th 1600 superpixels, as shown in Fig. 2C and 2D. Then the grey value of a superpixel is defined as the superpixel by *x*_i_, and its grey value by *I*(*x*_i_), with *i*=1,2,..,*N* and *N* = 1600.

**Figure 2.**
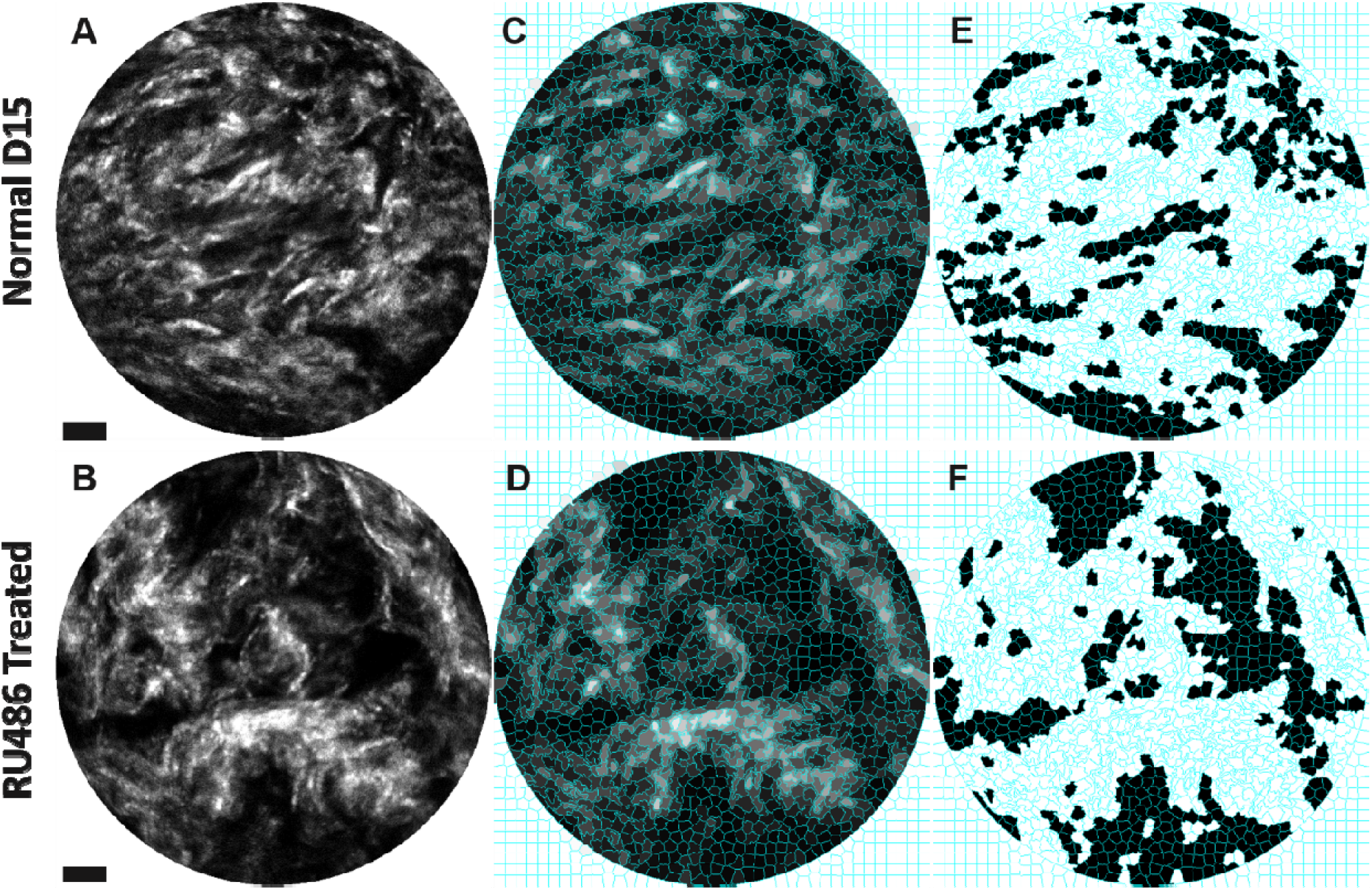
Demonstration of the superpixel-based graph-cut image segmentation. The two examples shown correspond to control group (top row) and RU486-treated group (bottom row), respectively. **A, B**, the cervical SHG images with pixel grey value proportional to the intensity of SHG signal. **C, D**, the corresponding superpixel images with boundaries of superpixels delineated in cyan. The grey value of each superpixel is assigned to be the average grey value of all constituent raw pixels. Note that the void margin outside the circular field of view are treated as collagen with saturated grey value. **E, F**, the segmentation result with *pore*-labeled superpixels colored in black and *collagen*-labeled ones in white.

Treating each superpixel as a graph node, an undirected graph *G*= (*V,E*) can be established, where edges in *E* connect spatially neighboring superpixels *V*={*x*_1,_*x*_2,…,_*x*_N_}in. Then the image segmentation task is essentially splitting the graph nodes (i.e. superpixels) into two classes, or equivalently, finding an optimal assignment of labels (*pore* or *collagen*) to all superpixels, by minimizing the following cost function:

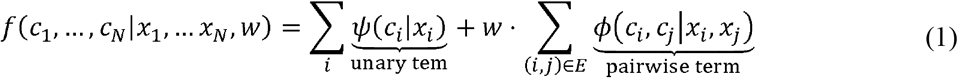

Where *c*_*i*_ ∈ {*pore, collagen*} is the class label assigned to superpixel *x*_i_ *ψ*(*c*_*i*_|*x*_i_) the unary cost (or penalty) of labeling the i-th superpixel as class *c*_*i*_ (as detailed shortly),ϕ(*c*_*i*_,-,*c*_*j*_,|*x*_*i*_,-,*x*_*j*_) the pairwise cost associated with labeling the *i-*th superpixel as class *c*_*i*_ and the *j-*th superpixel as class *c*_*j*_, respectively (as detailed shortly), and *w* regulates the relative significance between unary and pairwise cost terms.

The unary term *ψ*(*c*_*i*_|*x*_*i*_) assigns penalty to the *i*-th superpixel based on its class label *c*_*i*_ and grey value *I*(*x*_*i*_) In our implementation, this unary penalty term is defined as

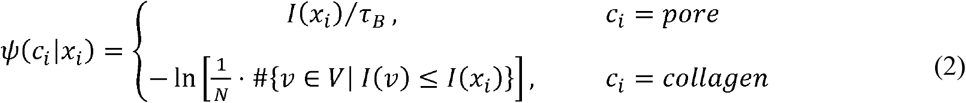

Here *τ*_*B*_ denotes the average grey-value level of background area, and #{*v ∈ V* | *I*(*v*) *≤ I*(*x*_*i*_)} denotes the number of superpixels in *V* with grey value no more than that of *x*_*i*._ According to Eq. (2), the penalty of labeling a superpixel as *pore* grows proportionally with its grey value *I*(*x*_*i*_), while the cost of labeling it as *collagen* decays with *I*(*x*_*i*_). Intuitively, the unary term inflicts penalty based on how close a superpixel’s grey value matches the expected grey value corresponding to class it is labeled; to be more specific, superpixels labeled as *collagen* tend to have larger grey value, while darker superpixels correspond more likely to the *pore* class.

Another operating principle for segmentation is that neighboring superpixels of similar grey values are more probable to carry identical labels. This intuition is taken care of in the cost function via the pairwise term, which is defined in our implementation as:

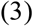

where is a normalization factor for inter-node difference in grey value. The pairwise term penalizes only those edges connecting two nodes of different classes, and for each such edge, the penalty itself diminishes exponentially with increasing grey value discrepancy between the two nodes it bridges. Such design therefore favors partitioning neighboring superpixels with abrupt change in grey value.

Finally, the relative significance between intra-node penalty (i.e. the unary term) and inter-node penalty (i.e. the pairwise term) is regulated via the weight factor *w*. Putting everything together, the optimization problem in Eq. (1) can be efficiently solved using the classical min-cut/max-flow graph-cut algorithm^36^.

In our implementation, with the grey value of raw pixels and superpixels ranging from 0 to 255, we determined the average background level *τ*_*B*_= 24 based on the grey value distribution of visually recognized background (super)pixels. The normalization factor for inter-node grey value difference *σ*_*E*_= 2 and regulation factor, *w*= 100 were decided via iterations of test and adjustment. Examples of final segmented superpixel-comprising images are shown in Fig. 2E and 2F. Within each segmented image, neighboring *pore-*labeled superpixels are merged into a single larger pore. Then the total number of pores are counted for each image, with the size of each pore given by summing the size of all constituent superpixels. The porosity fraction of each FOV can be calculated by dividing total pore area (sum of area of all pores) by the area of the circular FOV (100 µm in diameter).

### 2.5 Statistics

A total of ten cervical tissue sections of the control group (randomly selected from 5 normal pregnant mice at gestation day 15) and 16 cervical tissue sections of the treated group (randomly selected from 8 RU486-treated mice) were imaged; for each tissue section, ∼40-60 SHG images, each corresponding to an FOV ∼100 µm in diameter were acquired. Fiber diameter and porosity analysis were then performed on each SHG image (i.e. FOV). For each animal, the mean fiber diameter (i.e. the fitted 2*σ*. value), mean number of pores per FOV, and mean porosity fraction are simply given by the average of corresponding quantities over all images acquired from (all tissue sections of) the respective animal. The mean pore size, however, is calculated by averaging over the collection of all pores extracted from all images acquired from (all tissue sections of) the respective animal. With all these animal-wise mean values (n = 5 for day 15, and n = 8 for RU486 group), the average and standard deviation of mean values are calculated, and unpaired two-tailed Welch’s *t*-tests were performed in MATLAB (MathWorks, Natick, Massachusetts).

## 3. Results

Shown in Fig. 3 are representative collagen SHG images of cervical tissue sections of normal mice and PTB mouse models acquired using our newly-developed SHG endomicroscope. The overall difference in collage matrix morphology between normal pregnant mice at gestation day 15 (Figs. 3A-3C) and the RU486 PTB mouse models (Figs. 3D-3F) is stark. Specifically, cervical collagen fibers of the control group are distributed more uniformly in the corresponding SHG images, while the cervical collagen matrices of the RU486-treated group exhibit substantially larger pores. Those pore areas, void of SHG signal, could be tissue spaces occupied by cells, blood vessels, fluid (?), or extracellular matrix devoid of type I collagen. The enlarged inter-collagen spacing revealed by SHG images of RU486-treated cervical tissue sections reflects the dramatic collagen remodeling induced by mifepristone^37,38^.

**Figure 3.**
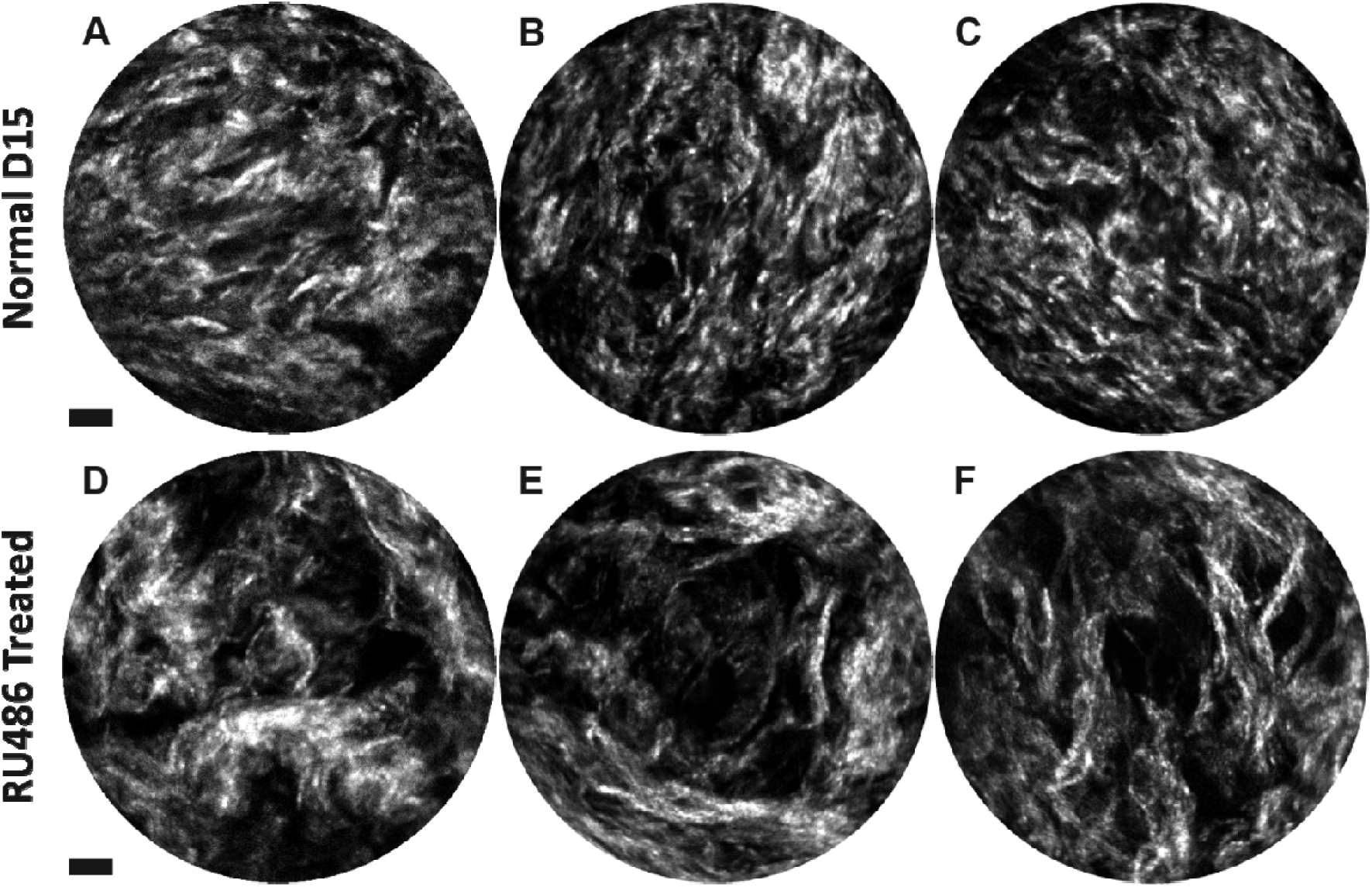
Representative endomicroscopic SHG images of cervical tissue sections. **A-C**, normal pregnant mice at gestation day 15. **D-F**, RU486 preterm birth mouse models. Ten-frames are averaged for the images here, corresponding to an equivalent acquisition time of ∼3.8 sec. Scale bar: 10 µm.

Quantitative characterization of collagen matrix morphology, including fiber diameter and porosity analysis, was undertaken as described in Section 2.3-2.5, with results illustrated in Fig. 4. The mean collagen fiber diameter of RU486-treated group is relatively larger than that of normal group (Fig. 4A, *p-* value < 0.005); cervices of treated mice also display on average larger mean pore size and a smaller number of pores (Figs. 4B and 4C, *p*-value < 0.005), but no significant change in terms of the overall porosity area fraction (Fig. 4C, *p*-value = 0.095). These quantitative results imply that mifepristone treatment does not seem to change the total cervical collagen content, but mainly alters the macroscopic organization of fibrillary collagen. The increase in pore size and fiber diameter, and the decrease in pore number, suggest that collagen fibers amalgamate with each other, forming relatively thicker fiber bundles and leaving fewer but enlarged collagen-void spaces (i.e. pores).

**Figure 4.**
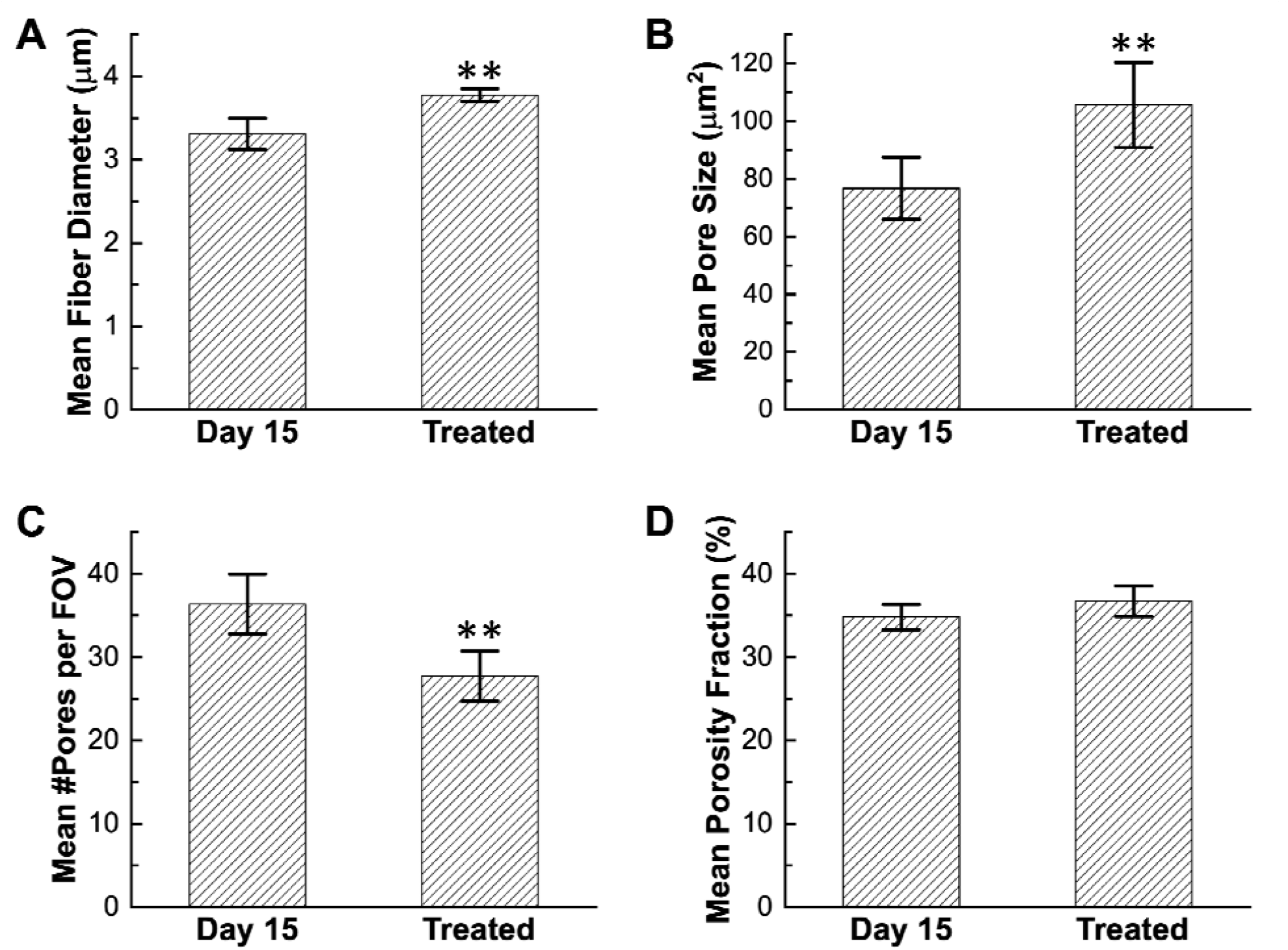
Fiber diameter and porosity analysis of cervical collagen. The average of mean values are compared between the control and RU486 treated group for: (A) fiber diameter scale fitted from the autocorrelation function, (B) individual pore size and (C) number of pores within each FOV, and (D) porosity fraction. Error bars represent standard deviation of the mean values over n = 5 control animals at gestation day 15, and over n = 8 animals from the mifepristone-treated PTB group, respectively. denotes *p*-value < 0.005.

To evaluate the applicability of our SHG endomicroscope in practical *in situ* imaging setup, we further imaged intact cervices directly (i.e., without sectioning). As described in Section 2.2, the endomicroscope needs to image through the intact ectocervical epithelium to access collagen inside. Thanks to the 200 µm working distance afforded by the miniature objective on the current endomicroscope, the excitation laser can successfully penetrate deep enough to reach the subepithelial collagen network. Representative SHG images acquired in this fashion are shown in Fig. 5, from which one can clearly see that subepithelial collagen network of the mifepristone-treated group manifests generally increased porosity compared with the control group. The trend of changing in collagen porosity captured from intact cervices directly aligns well with that observed on cervical tissue sections as shown in Fig. 3.

**Figure 5.**
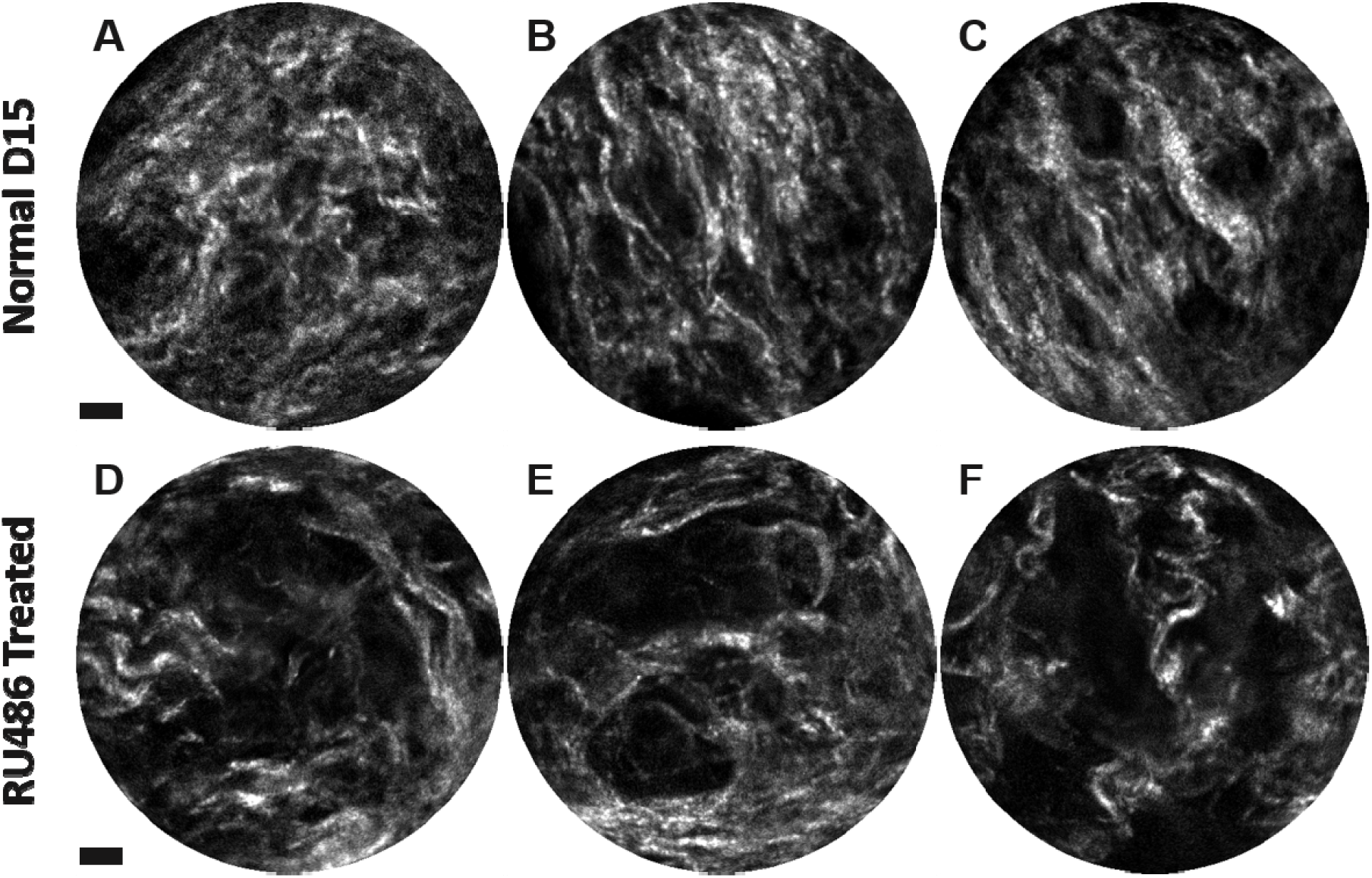
Representative SHG endomicroscopy images of sub-epithelial cervical collagen acquired directly from intact mouse cervices. Top row (A-C): normal pregnant mice at gestation day 15. Bottom row (D-F): RU486 preterm birth mouse models. In general, images of the treated group show increased porosity. Scale bar: 10 μm.

## 4. Discussion

The axial resolution of the SHG endomicroscope used in this study is ∼6.5 µm, which is about three times as thick as the axial resolution (typically ∼2 µm) of a bench-top SHG microscope^39^. Consequently, each SHG image acquired by our SHG endomicroscope equates to the axial accumulation of three layers of bench-top SHG microscopy images. As pore regions in the middle layer might get “contaminated” by collagen SHG signal leaking from the layer(s) above and/or below, the contrast between collagen-rich and collagen-void pore regions in endomicroscopy SHG images is less salient than in microscopy images. Due to such reduced collagen-to-pore contrast, the conventional porosity analysis approach which involves global threshold-based image segmentation followed by particle analysis^18,21,33^ doesn’t work well on our endomicroscopy SHG images, since there exists no global threshold that could accurately separate collagen from pore regions throughout the entire image. To overcome this limitation, we designed the superpixel-based graph-cut segmentation algorithm which accounts for both intensity and spatial proximity information of the superpixels and performs context-sensitive image segmentation. As demonstrated in Section 2.4 and Fig. 4, our algorithm can capture the porosity properties of endomicroscopic SHG images robustly and accurately, yielding statistical insights of cervical collagen morphology that are otherwise unavailable using the conventional global threshold-based segmentation.

Previous SHG microscopy-based study has reported similar increase in porosity of cervical collagen matrix in response to mifepristone treatment, demonstrating decreased number of pores and increased pore sizes^18,38^. One different observation herein is that the endomicroscopy SHG images of cervical collagen from mifepristone-treated group manifest no significant increase in total pore area fraction. Such discrepancy could stem still from the inferior axial resolution, which leads to that each epi-SHG image acquired by our endomicroscope spans an axial layer three times as thick as that acquired by a benchtop SHG microscope and therefore contains richer SHG content and less pore area fraction. To further investigate this hypothesis, one potential direction is to acquire with a benchtop high-NA SHG microscope axially densely-sampled three-dimensional (3D) volumetric image stacks and then quantify the 3D collagen matrix morphology across all voxels within each SHG image volume^40^.

The increase in porosity of the subepithelial collagen matrix of intact mifepristone-treated cervices (Fig. 5), although conspicuous in quite a few endomicroscopic SHG images, was not consistently seen across all imaging locations over the ectocervical epithelium. In contrast, tissue sections collected throughout the longitudinal extent of mifepristone-treated cervices show consistently higher porosity than those from the control group (Fig. 3 and Fig. 4). Given the 200-μm working distance of our SHG endomicroscope, the lack of uniform increase in porosity of subepithelial SHG images acquired from intact mifepristone-treated mouse cervices implies that the collagen matrix residing immediately beneath the ectocervical epithelium could have undergone less dramatic remodeling, and therefore doesn’t exhibit as pronounced porosity increase as stromal collagen matrix deeper beneath the ectocervical epithelium. Towards the overarching goal of non-invasive diagnostic imaging, further technology development towards deeper penetration as well as enlarged FOV size is needed for SHG imaging to extract consistent, statistically significant changes in cervical collagen morphology (including but not limited to porosity)^41^.

In summary, we have developed a high-performance, flexible miniature SHG endomicroscope which enables label-free, non-invasive profiling of cervical collagen morphology, and a tailored superpixel-based graph-cut segmentation algorithm for quantitative porosity analysis of endomicroscopic SHG images. The significant increase in collagen porosity revealed in endomicroscopic SHG images of cervical tissue sections from mifepristone-treated mouse models promises promising possibilities for non-invasive imaging-based early diagnosis and risk assessment of preterm birth *in vivo*.

